# Transfer of Antibiotic Resistance Genes from Gram-positive Bacterium to Gram-negative Bacterium

**DOI:** 10.1101/2020.11.01.364331

**Authors:** Prasanth Manohar, Thamaraiselvan Shanthini, Bulent Bozdogan, Cecilia Stalsby Lundborg, Ashok J Tamhankar, Nades Palaniyar, Nachimuthu Ramesh

## Abstract

The emergence of antibiotic resistance due to uncontrolled use of antibiotics in non-humans, poses a major threat for treating bacterial infections in humans. Added to this is the possibility of transfer of resistance from Gram-positive bacteria to Gram-negative bacteria. Therefore, the possibility of resistance gene transfer from a non-human originated pathogenic bacterium to a pathogenic bacterium infecting humans needs evaluation. In this study, poultry litter samples collected from Tamil Nadu, India were screened for the presence of meropenem- and cefotaxime-resistant *Staphylococcus sciuri*. Standard microbiological techniques and 16S rRNA analysis were used to confirm *S. sciuri*. In the resistant isolates, resistance genes such as *bla*_NDM-1_, *bla*_OXA-48-like_, *bla*_KPC_, *bla*_VIM_, *bla*_IMP_ and *bla*_CTX-M_ were screened. Transconjugation studies were performed using donor, *S. sciuri* and recipient, *E. coli* AB1157 (Str^r^). A total of 26 meropenem-resistant and 24 cefotaxime resistant *S. sciuri* were isolated from poultry litter samples. The presence of *bla*_NDM-1_ (n=2), *bla*_IMP_ (n=8), *bla*_CTX-M-9_ (n=5) and *bla*_CTX-M-2_ (n=1) was detected. Transconjugation results confirmed that *S. sciuri* carrying plasmid-borne resistance gene *bla*_NDM-1_ conjugated to *E. coli* AB1157. The transferability of resistance genes from *S. sciuri* to *E. coli* could be another possible reason for spread of antibiotic-resistant bacteria in humans.

## 1. Introduction

Antibiotic resistance due to uncontrolled use of antibiotics poses a major threat to treat bacterial infections in humans and WHO has described antibiotic resistance as the greatest malice of 21^st^ century [1]. The use of antibiotics in animals is linked to development of drug-resistant bacterial infections in humans and animals [2]. In countries such as the US and in the European countries such as the UK, Sweden and the Netherlands, necessary steps have been taken to control the use of antibiotics in animals [3-5]. But in India, multiple antibiotics in large amounts are routinely used in farm animals, especially in poultry industry [6-8]. Due to this practice, selection for resistance in natural microbiota on/in animals and those in their environment has grown progressively [9]. The antibiotic resistant bacteria from animals can potentially enter the human bacterial populations through various pathways such as through direct contact, animal meat and farming practices [10-13]. Bacteria carrying resistance genes in extra-chromosomal DNA (r-plasmid) can be shed in the feces, from where these genes can be disseminated to other bacterial pathogens. Accumulation of resistant bacteria in animals can result in the spread of antibiotic resistant bacteria in humans [14]. Concerns have therefore been raised against the use of antibiotics in animal feed, as this practice could contribute to the spread of antibiotic resistant bacteria in humans [15].

The antibiotics used in livestock can enter the environment; for example, many antibiotics are excreted unchanged and when such excreta is used as manure, they reach the environment through manure application and in to agro-ecosystem receiving the agricultural waste. Use of antibiotics in livestock also creates the potential for antibiotic resistance generation and transmission through the food processing chain [16]. In India, the use of antibiotics in food animals is high compared to other countries (both the quantity of animals treated as well as concentrations of antibiotics are high), with poultry industry being the highest user [6, 17]. There are no regulatory guidelines in India for the use of antibiotics in chicken, cattle and pigs that are raised for domestic consumption. Previous studies in India also showed the presence of antibiotic residues in food animal products (chicken meat), indicating the widespread of antibiotic use in food animals [6, 18,19].A study showed that residues of five antibiotics were found in the chicken tissues ranging between 3.37-131.75 μg/kg which is above the permissible maximum residue limit (MBL) of 100 μg/kg [20]. Of the 40 per cent samples found tainted with antibiotic residues, 22.9 per cent contained residues of only one antibiotic while the remaining 17.1 per cent samples had residues of more than one antibiotic. This is the biggest study done in India to test residues of antibiotics in chicken [17]. Another study conducted by the Center for Disease Dynamics, Economics and Policy (CDDEP) in Punjab, (India) revealed that the poultry samples collected from the farms that were using antibiotics had three-times higher multi-drug resistant bacteria than the farms that did not use antibiotics for growth promotion [21]. Further, poultry litter has also been found to carry antibiotic resistant bacteria and resistance associated genes [22].

*Staphylococcus sciuri* is a Gram-positive bacterium and one of the common florae found in poultry. It has been reported that coagulase negative staphylococci such as *S. sciuri* can colonize poultry intestine and can cause infections [23,24]. *S. sciuri* can cause infections in humans also and it has been reported that 0.79-4.3% of coagulase-negative *Staphylococci* isolated from (human) clinical samples were *S. sciuri* [25]. Instances of their outbreaks in hospitals have been reported [26-28]. *Escherichia coli*, a Gram-negative bacterium, can cause community and hospital-acquired infections in humans including diarrheal diseases. *E. coli* forms part of the bacterial flora of the human gastrointestinal tract, and also acts as a reservoir for harboring antibiotic resistance genes [29]. Transfer of plasmid carrying antibiotic resistance genes from Gram-positive bacterium to *E. coli* has been reported [30]. Thus, the possibility of gastrointestinal colonization by *E. coli* which has acquired resistance from *S. sciuri* through horizontal gene transfer through a plasmid carrying a resistance gene cannot be totally ruled out.

We have undertaken a project to determine the antibiotic resistance profile of bacteria isolated from poultry litter and the potential risk they pose in disseminating antibiotic resistance through plasmids. Though we have isolated several Gram-positive and Gram-negative bacteria from poultry litter, this paper describes the isolation of Gram-positive bacterium *S. sciuri* from poultry litter, its antibiotic resistance profile, and its potential to transconjugate its plasmid-borne resistance gene to Gram-negative bacterium, *E. coli*.

## 2. Materials and Methods

### 2.1 Study setting and sampling procedure

The southern part of Tamil Nadu, India, is one of the most highly concentrated areas of poultry production with ≥100 poultry farms within an area of about 100 km^2^ and each farm having the capacity of 10,000 birds (14,000 sq.ft). It is also an area with high agricultural production, and most of the poultry waste from these farms is applied as manure for agricultural lands, increasing the possibility of human-poultry litter contact as in India most of the agricultural operations are conducted using human labour and not machines. We collected poultry litter samples from three poultry farms (Farms A, B and C) separated by a distance of 15 km each. In this area, the chickens remain on the farms for a period of about 85 days before sending them for processing. Therefore, we collected the poultry litter samples at ∼85 days to ensure that the birds have received the typical drug regimens (Table 1). One sample (≈25g) was collected from each poultry farm and the poultry litter was collected from different locations within the same sampling farm (no statistical method was used). The samples (n=3) were collected in sterile containers (Himedia, India) and immediately stored at 4°C. All the samples were transported for antibiotic susceptibility testing to Antibiotic Resistance and Phage Therapy Laboratory (VIT) within 10 hours of collection under cold conditions (4°C).

**Table 1:** Antibiotics used in poultry farms from which litter samples were collected.

| Antibiotics and purpose of use | Farm A | Farm B | Farm C |
| --- | --- | --- | --- |
| **Amoxicillin:** For bacterial infections. (with drinking water) | - | 1 mg/5mL, starting from day 3 to 6<sup>th</sup> week. | - |
| **Bacitracin:** For growth promotion (with feed) | 20 mg/kg b.w/day for 5 days. | 20 mg/kg b.w/day for 5 days. | 15 mg/kg b.w/day for 5 days. |
| **Ciprofloxacin:** To treat respiratory and gastrointestinal tract infections.(with drinking water) | 1 mg/mL, starting from day 3 to 6<sup>th</sup> week. | 1 mg/5mL, starting from day 3 to 6<sup>th</sup> week. | 1 mg/5mL, starting from day 3 to 6<sup>th</sup> week. |
| **Chlortetracycline:** For bacterial infections (prophylaxis).(with drinking water) | 250 mg/L for 8 days from onset of infection. | - | 500 mg/L for 8 days. |
| **Chlortetracycline:** For growth promotion(with feed) | 20 mg/kg b.w/day for 5 days. | 20 mg/kg b.w/day for 5 days. | 15 mg/kg b.w/day for 5 days. |
| **Erythromycin:** To prevent infections caused by micro-organisms especially cocci. (with drinking water) | - | 0.5 g/L for 5-8 days from onset of infection. | - |
| **Erythromycin:** For growth promotion(with feed) | 20 mg/kg b.w/day for 5 days. | - | 15 mg/kg b.w/day for 5 days. |
| **Gentamicin:** To control colibacillosis and paratyphus at early stage(subcutaneous). Dosage depends on growth. | 0.2 mg/0.2 mL for each-day-old chickens | 1 mg/mL dose for each-day-old chickens. | 0.5 mg/mL dose for each-day-old chickens. |
| **Linomycin:** For growth promotion (with feed) | 20 mg/kg b.w/day for 5 days. | 20 mg/kg b.w/day for 5 days. | - |
| **Neomycin:** Effective against bacterial infections. (with drinking water) | - | - | 1 mg/5mL, starting from day 3 to 6<sup>th</sup> week. |
| **Neomycin:** To increase weight. (with feed) | - | 20 mg/kg b.w/day for 5 days. | - |
| **Oxytetracycline:** To prevent infections caused by *Mycoplasma* and *E. coli*.(with drinking water) | 1000 mg/gallon for 15 days from onset of infection. | 800 mg/gallon for 15 days from onset of infection. | - |
| **Penicillin:** For growth promotion(with feed) | 20 mg/kg b.w/day for 5 days. | 20 mg/kg b.w/day for 5 days. | 15 mg/kg b.w/day for 5 days. |
| **Streptomycin:** To treat enteric diseases caused by coliform bacteria. (with drinking water) | - | - | 1 mg/5mL, starting from day 3 to 6<sup>th</sup> week. |
| **Virginiamycin:** For growth promotion (with feed) | 20 mg/kg b.w/day for 5 days. | 20 mg/kg b.w/day for 5 days. | - |

Data provided in table was obtained from the health professionals taking care of the birds in respective poultry farms. b.w.-body weight; “-“ means not used.

### 2.2 Isolation of *S. sciuri* from poultry litter

Samples from the three different poultry farms (one sample from each farm) were separately serially diluted and resistant bacteria were isolated. Briefly, 5 g of each sample was diluted in 45 mL of 0.1% peptone water, and 1 mL from this mixture was serially diluted (10^−1^ to 10^−8^) in 0.1% peptone water, as described previously [31]. From each serially diluted tube (10^−8^ dilution representing each poultry farm), 100 µL was taken and plated on to Mueller Hinton agar plates that were supplemented with antibiotics (meropenem and cefotaxime) based on their break point values (meropenem ≥4 mg/L, cefotaxime ≥4 mg/L) according to CLSI guidelines, 2016 [32]. Plates were incubated for 18 hours and colony forming units (CFU) were calculated. Though there was appearance of numerous bacteria, *S. sciuri* was chosen based on the morphological characteristics of diffusible pigment production, and yellowish-grey colony and sub-cultured thrice with same concentration of antibiotics from which it was isolated. Resistant isolates were stored in 30% glycerol with brain heart infusion medium at −20°C until screened further for genomic characteristics.

### 2.3 Identification of *S. sciuri*

During the isolation process, any bacterial colony that resembled staphylococci was isolated and further subjected to identification procedure. Preliminary identification of an isolate to be *S. sciuri* was based on microscopic characterization, positive oxidase test, positive catalase test and use of VITEK 2 identification system (bioMerieux). DNA isolation was performed for bacterial isolates using HiPurA Bacterial genomic DNA purification Kit (Himedia, India). Species amplification of 16S rRNA gene from each isolate was done and sequenced (Eurofins Genomics, India). Universal primer sequences were used for 16S rRNA amplification; 27F: 5’-AGAGTTTGATCMTGGCTCAG-3’ and 1492R: 5’-CGGTTACCTTGTTACGACTT-3’, for phylogenetic study as they are highly conserved between different species of bacteria. Sequence identities were determined using nucleotide blast at www.ncbi.nlm.nih.gov/BLAST/ and species level identification was made. In BLAST, nucleotide sequence similarity search was performed, and sequences having an identity of 98% and above were considered as sufficient for species identification. Sequences were submitted to NCBI “Genbank” and accession numbers were obtained.

### 2.4 Screening of antibiotic resistance genes

All the resistant (meropenem and cefotaxime) *S. sciuri* isolates (n=50) were chosen for gene amplification. Accordingly, for meropenem resistant isolates *bla*_NDM-1_, *bla*_OXA-48-like_, *bla*_IMP_, *bla*_VIM_ and *bla*_KPC_ carbapenemase genes were screened by multiplex PCR using primers and reaction conditions described by Doyle *et al*., [33], and cefotaxime resistant isolates were amplified with *bla*_CTX-M_ group genes by multiplex PCR using conditions and primers described by Mirzae*et al*., [34]. Primers used for PCR amplification are presented in Table S1.

### 2.5 Plasmid profiling and transconjugation study

Plasmid DNA was isolated independently for all the isolates carrying resistance gene. Bacterial cultures were grown for 16 hours and plasmids were isolated using HiPurA plasmid DNA miniprep purification kit as per manufacturer’s instructions (Himedia, India). Isolated plasmids were loaded on to agarose gels to confirm the presence of plasmids in each bacterial isolate and lambda DNA digested with EcoRI and HindIII was used as a marker. The isolated plasmids were used as a template for PCR reactions to screen antibiotic resistance genes.

Conjugation studies were carried out using the double selection method [35]. Transconjugation experiments were performed using *E. coli* AB1157 (Str^r^) as a recipient [36]. For transconjugation experiments, the isolate that carried resistance genes in their plasmid DNA (serving as a donor) was selected and grown overnight in MH broth containing meropenem and *E. coli* AB1157 in medium with streptomycin. Donor and recipient bacteria were mixed together at a 9:1 ratio (v/v) for the maximum availability of donors for the recipient [36]. The mixture was kept undisturbed for 4 hours at 37°C. From this; 0.1 mL was then spread on to MH agar plates containing both meropenem (8 mg/L) and streptomycin (100 mg/L). The control plates included donor isolate spread plated on MH agar plates containing meropenem (8 mg/L), donor isolate spread plated on MH agar plates containing streptomycin (100 mg/L), *E. coli* AB1157 spread plated on MH agar plates containing meropenem (8 mg/L) and *E. coli* AB1157 spread plated on MH agar plates containing streptomycin (100 mg/L). The experiment was repeated in triplicate. Efficiency of conjugation was determined by the number of transconjugants per donor cell and genotypically through amplification of *bla*_NDM-1_ in transconjugants.

## 3. Results

### 3.1 Antibiotic resistant *S. sciuri* bacteria are present in poultry reared in all the three farms that routinely use antibiotics

To determine whether antibiotic use for poultry production promotes antibiotic resistance in *S. sciuri*, we collected poultry litter from three farms that routinely use multiple antibiotics and analyzed the *S. sciuri* isolated from these samples. These farms administered many antibiotics (Amoxicillin, Bacitracin, Chlortetracycline, Ciprofloxacin, Erythromycin, Gentamicin, Linomycin, Neomycin, Oleandomycin, Oxytetracycline, Penicillin, Streptomycin, Virginiamycin) via various routes (subcutaneously, in drinking water and with feed). Although the antibiotic regimens were slightly different, all the three farms used multiple antibiotics for several days (Table 1). The confirmation of *S. sciuri* isolates was based on colony morphology (size of the bacterial colony 3-4 mm), Gram staining, microscopic characterization, positive oxidase test, positive catalase test and VITEK 2 identification system (bioMerieux). A total of 50 non-repetitive, meropenem- and cefotaxime-resistant *S. sciuri* were isolated from poultry litter samples. These isolates included 26 meropenem resistant *S. sciuri*: 10 from Farm A, 8 from Farm B and 8 from Farm C, and 24 cefotaxime resistant *S. sciuri*: 9 from Farm A, 7 from Farm B and 8 from Farm C (Table 2, S2). 16S rRNA analysis confirmed that the isolates were *S. sciuri*. These sequences have been deposited in Genbank (Sequences KT779211-KT779218; https://www.ncbi.nlm.nih.gov/nuccore/). These isolates were distributed among the 3 farms without any specific predominance suggesting that antibiotic resistant *S. sciuri* were generally distributed on all the farms and antibiotic use on the farms may be generating antibiotic resistance.

**Table 2:** The distribution of resistance genes among meropenem- and cefotaxime-resistant *S. sciuri* isolated from poultry litter.

| Poultry farm | Meropenem-resistant *S. sciuri* | Presence of *bla*<sub>NDM-1</sub> | Presence of *bla*<sub>IMP</sub> | Cefotaxime-resistant *S. sciuri* | Presence of *bla*<sub>CTX-M-9</sub> | Presence of *bla*<sub>CTX-M-2</sub> |
| --- | --- | --- | --- | --- | --- | --- |
| Farm A | 10 | 2 | 2 | 9 | 3 | 0 |
| Farm B | 8 | 0 | 3 | 7 | 1 | 1 |
| Farm C | 8 | 0 | 3 | 8 | 1 | 0 |

### 3.2 Prevalence of antibiotic resistance genes in *S. sciuri*

When the resistance genes for each antibiotic class were amplified, amplification results showed that out of 26 phenotypically meropenem resistant isolates (*Staphylococcus sciuri* meropenem-SCM), only 2 had *bla*_NDM-1_ (SCM3, SCM7) and 8 had *bla*_IMP_ (SCM5, SCM10, SCM13, SCM15, SCM18, SCM19, SCM21, SCM25) and none had *bla*_OXA-48-like_, *bla*_VIM_ or *bla*_KPC_. Among 24 cefotaxime resistant isolates (*Staphylococcus sciuri* cefotaxime-SCC), 5 had *bla*_CTX-M-9_ (SCC29, SCC31, SCC35, SCC39, SCC44) and 1 had *bla*_CTX-M-2_ (SCC37) gene (Table 2). The remaining isolates did not amplify for any of the above screened resistance genes. When 16 *S. sciuri* isolates that carried resistance genes were examined for the presence of plasmids, one isolate (*S. sciuri* SCM3 isolated from Farm A) carrying two different sized plasmids (bands) was detected (Fig.1). The isolated plasmid was screened for respective resistance genes and it was found that one isolate (*S. sciuri* SCM3) harboured *bla*_NDM-1_ gene in its plasmid DNA.

**Figure 1.**
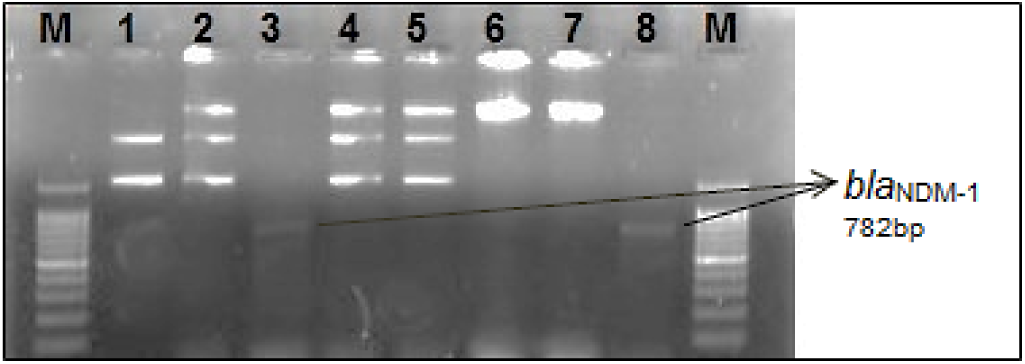
Confirmation of transconjugants carrying conjugative plasmids from the donor, ***S. sciuri***. Lane M: 100 bp DNA marker, Lane 1: Plasmids isolated from donor (*S. sciuri*), Lane 2: Total DNA with plasmids isolated from donor, *S. sciuri* (1:1 mixture of plasmid and DNA), Lane 3: Amplified *bla*_NDM-1_ from the plasmid isolated from donor (*S. sciuri*), Lanes 4-5: total DNA with plasmids isolated from transconjugate, Lane 6: Total DNA isolated from recipient (*E. coli* AB1157), Lane 7: Total DNA isolated from donor (*S. sciuri*), Lane 8: Amplified *bla*_NDM-1_ from the plasmid isolated from transconjugant. The amplified *bla*_NDM-1_ in the plasmid isolated from donor (*S. sciuri*) is same as the *bla*_NDM-1_ isolated from the transconjugant plasmid.

### 3.3 Transconjugation studies

To determine the ability of the transfer of antibiotic resistance genes from *S. sciuri* to *E. coli*, a plasmid-based transfer assay or transconjugation assay was conducted. When plasmid-borne *bla*_NDM-1_ from *S. sciuri* SCM3 (donor) was transformed to recipient strain (*E. coli* AB1157), the new *S. sciuri* plasmid was detectable in *E. coli* (donor to recipient). These transconjugants were examined for the presence of conjugative plasmid and the recipient strain, *E. coli* AB1157 had acquired meropenem resistance, which was confirmed genotypically by amplification of *bla*_NDM-1_ gene (Fig.2). Conjugation efficiency was calculated to be 8.35 × 10^−5^ (Table 3). Therefore, it was confirmed that antibiotic resistance gene from *S. sciuri* is transferable to *E. coli* through conjugative plasmid.

**Figure 2.**
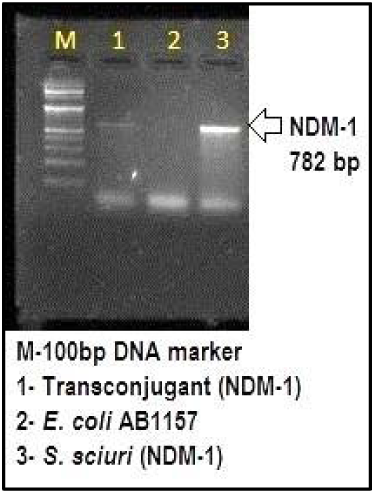
Comparison showing transconjugants and donor *S. sciuri* carrying NDM-1 gene in agarose gel electrophoresis. M – 100bp ladder; 1 –PCR product of NDM-1 gene amplified from plasmids isolated from transconjugants; 2 - *E. coli* AB1157 DNA carrying no gene; 3 - PCR product of NDM-1 gene amplified from plasmids isolated from donor *S. sciuri*.

**Table 3.** Conjugation efficiency of plasmids carrying *bla*_NDM-1_ from *S. sciuri* to *E. coli*. Plasmid-borne *bla*_NDM-1_ gene from donor *S. sciuri* was used for transconjugation experiments with recipient *E. coli* AB1157.

| Donor used for conjugation | Number of donors in 1 mL | Number of recipients in 1 mL | Donor : recipient ratio | Number of transconjugants in 1 mL | Efficiency of conjugation* |
| --- | --- | --- | --- | --- | --- |
| *S. sciuri* SCM3-*ndm-1* | $1.46 \times 10^9$ | $2.31 \times 10^9$ | 9:1 | $1.22 \times 10^5$ | $8.35 \times 10^{-5}$ |

Following conjugation, total viable recipients were selected for by plating on MH plates containing streptomycin. Transconjugants were selected for on MH plates containing both streptomycin and meropenem. \*Efficiency of conjugation is estimated as the number of transconjugants per donor cell.

## 4. Discussion

Emergence of antibiotic resistance due to routine use of antibiotics in poultry industry is a major concern, and the extent and role of antibiotic resistance in *S. sciuri* in this context is not well established. Transfer of antibiotic resistance gene between Gram-positive and Gram-negative bacteria was reported previously [30, 37, 38]. This study showed that *S. sciuri* isolated from poultry litter samples from all the poultry farms, possessed resistance to meropenem (one of the last resort antibiotics) and cefotaxime (extended-spectrum beta-lactamase antibiotic). While resistance genes *bla*_NDM-1_, *bla*_IMP_, *bla*_CTX-M-9_ and *bla*_CTX-M-2_ were detected, *bla*_OXA-48-like_, *bla*_VIM_, *bla*_KPC_, *bla*_CTX-M-1_, *bla*_CTX-M-8_ and *bla*_CTX-M-25/26_ were not detected. Plasmid profiling showed the presence of conjugative elements. Transconjugation studies performed using *S. sciuri* as donor and *E. coli* AB1157 as recipient, showed a conjugative efficiency of 8.3 × 10^−5^ for *bla*_NDM-1_ positive staphylococcal isolate. This study highlights the emergence of mono- and multi-drug resistance in *S. sciuri* and its ability to transfer the resistance gene to *E. coli* through plasmid transfer.

Previous studies investigating bacteria from poultry litter samples suggested the presence of antibiotic resistant bacteria and its potential role in disseminating antibiotic resistance [31-33]; poultry litter from antibiotic-intensive farms could be an important source for the emergence and global spread of antibiotic resistant bacteria. In general, strains of *S. sciuri* under normal circumstances do not carry antibiotic resistance genes [39-41]. However, earlier studies have indicated that *S. sciuri* isolated from farm animals could harbour plasmids with antibiotic resistance genes (*cfr, optrA, fexA, aadD, erm*) [39, 40]. Horizontal gene transfer and conjugative plasmid transfer between a few Gram-positive and Gram-negative bacteria [42, 43], and transfer of plasmid (pKJK10) among indigenous bacteria in the barley rhizosphere was earlier demonstrated [43]. However, evidence for the possible horizontal transfer of plasmid-borne resistance genes from *S. sciuri* to *E. coli* or from animal borne organisms to those infecting humans is lacking. In this study, we report successful transfer of plasmid-borne *ndm-1* resistance gene from *S. sciuri* isolated from poultry litter to *E. coli*. As both *S. sciuri* and *E. coli* can colonize chicken intestine, causing gut-associated infections, there is possibility of resistance gene transfer *in vivo*. Avian pathogenic *E. coli* can cause local infections in chicken, such cellulitis, omphalitis, salpingitis and synovitis [44]. A study reported the presence of *cfr* genes among *Staphylococcus* sp. and *E. coli* isolates from the same farm [45]. This could be probably due to the spread of *cfr* across species and genus boundaries, mainly via horizontal gene transfer mediated by mobile genetic elements including plasmids [45]. To our knowledge, successful transconjugation experiments from *S. sciuri* to *E. coli* either do not exist or are very rare. We have reported here one such rare successful demonstration of the possibility of horizontal gene transfer from *S. sciuri* to *E. coli*.

Meropenem is considered as one of the last resort antibiotics for human infections [46]. Although they were not included in the food regimen for poultry in the farms from where the litter samples were taken for intercepting bacteria for resistance tests in this study, there was still a high degree of resistance observed to them. Out of 26 phenotypically meropenem resistant isolates, 2 had *bla*_NDM-1_ and 8 had *bla*_IMP_ gene. The *bla*_NDM-1_ gene was plasmid-borne, which indicated its potential for horizontal gene transfer. When experimentally transconjugated, the *bla*_NDM-1_ gene could be successfully transferred to *E. coli*, a bacteria responsible for several common infections and made it resistant to the concerned antibiotic. This sheds light on probably one of the mechanisms of resistance transfer, from Gram-positive to Gram-negative bacteria, and therefore on the reason for continuously increasing resistance to antibiotics that were earlier effective. The environmental aspect of generation and spread of antibiotic resistance and also the zoonotic aspect of it is still a neglected issue and there is an immediate need to focus on such studies [47]. Existence of genes with the potential for horizontal gene transfer (plasmid borne resistance) in *S. sciuri*, one of the bacteria commonly found in poultry litter, with which farmers come in contact in farming practice, strongly suggests that the uncontrolled use of antibiotics in poultry should be stopped to control the development and dissemination of antibiotic resistance, otherwise no last resort lifesaving antibiotics will remain available for human infections resulting in serious consequences for humanity. *S. sciuri* is associated with animals but its clinical relevance for humans has developed rapidly [22,48,49]. These staphylococci resistant to carbapenem and cephalosporin group of antibiotics should be monitored especially from animal husbandry and also the surrounding environmental niches. Dissemination of resistant genes *bla*_NDM-1_ or any other resistance genes to other bacteria (pathogens) through fecal waste can be controlled by pre-treatment steps to eliminate the bacterial load. Screened antibiotic resistant genes *bla*_NDM-1_, *bla*_IMP_ and *bla*_CTX-M_ from staphylococci were the most important part of this study because these highly prevalent resistance genes in human pathogens were screened from bacteria isolated from poultry litter. Most importantly, the identified resistance genes were found to have a potential for horizontal gene transfer.

Plasmid profiling also indicated the presence of antibiotic resistance gene in extra-chromosomal DNA of *S. sciuri* that had a capability to transfer resistance with high conjugation efficiency. The clustering of these resistance genes in conjugative plasmids can increase the persistence of antibiotic resistance [50]. This paper brings into focus bacteria carrying plasmid-encoded metallo-beta-lactamase carbapenem resistance genes in poultry environment, which carry a potential of horizontal gene transfer to other common pathogens that could pose a very serious threat to human medicine. To resolve such issues of transfer of genes from one environmental niche to another needs a ‘One Health approach’. The global action plan on antimicrobial resistance emphasizes the ‘One Health approach’ i.e. seeing humans, animals, the food chain, the environment, and the interconnectedness between them as one entity [51, 52]. Such ‘One Health’ studies are very important in countries like India [53], where the barriers to contacts between humans, animals and their environment are very weak.

## 5. Conclusions

*Staphylococcus sciuri*, a bacterium commonly associated with animals including poultry and increasingly being implicated in human infections, was found to carry extra-chromosomal, plasmid encoded resistance genes-*bla*_CTX-M_, *bla*_NDM-1_ and *bla*_IMP_, out of which in transconjugation experiment, plasmid-borne bla_NDM-1_ got transferred to a common pathogen *E. coli*. Such a horizontal gene transfer from *S. sciuri* to *E. coli* portents a dangerous potential threat in terms of spread of antibiotic resistance and shows poultry litter as a reservoir *o*f antibiotic resistance.

## Supplementary Materials

Table S1. Primer sequences used to amplify resistance genes in this study-*bla*_NDM-1,_ *bla*_OXA-48-like,_ *bla*_IMP_, *bla*_KPC_ and *bla*_VIM_ for metallo-beta lactamase and *bla*_CTX-M_ group gene for extended spectrum beta-lactamase. Table S2: Distribution, prevalence of resistance genes and conjugation studies using *S. sciuri* isolated from poultry litter.

## Author Contributions

Authors PM, TS and NR, collected the isolates and did resistance screening. Authors PM, TS, BB and NR undertook the laboratory work, AJT, CSL, NR and NP interpreted the data, and PM and NR wrote the initial manuscript. Authors AW, CSL, AJT and NP revised and edited the manuscript.

## Acknowledgments

The authors would like to thank Vellore Institute of Technology for providing the research facilities to complete the work.

