## Supplementary Table 1 for "Transfer of Antibiotic Resistance Genes from Gram-positive Bacterium to Gram-negative Bacterium"

**Supplemental information**

**Table S1. Primer sequences used to amplify resistance gene in this study.** *bla*_NDM-1,_ *bla*_OXA-48-like,_ *bla*_IMP_, *bla*_KPC_ and *bla*_VIM_ for metallo-beta lactamase and *bla*_CTX-M_ group gene for extended spectrum beta-lactamase.

| **GENE NAME** | **PRIMER SEQUENCE** | **REFERENCE** |
| --- | --- | --- |
| *bla*_NDM-1_ | F 5′-GCAGCTTGTCGGCCATGCGGGC-3′  R 5′-GGTCGCGAAGCTGAGCACCGCAT-3′ | 32 |
| *bla*_OXA-48-like_ | F 5′-GCGTGGTTAAGGATGAACAC-3′  R 5′-CATCAAGTTCAACCCAACCG-3′ | 32 |
| *bla*_IMP_ | F 5′-GAAGGCGTTTATGTTCATAC-3′  R 5′-GTACGTTTCAAGAGTGATGC-3 | 32 |
| *bla*_VIM_ | F 5′-GTTTGGTCGCATATCGCAAC-3′  R 5′-AATGCGCAGCACCAGGATAG-3′ | 32 |
| *bla*_KPC_ | F 5′-TGTCACTGTATCGCCGTC-3′  R 5′-CTCAGTGCTCTACAGAAAACC-3′ | 32 |
| CTX-M Group-1 | F 5′-AAAAATCACTGCGCCAGTTC -3′  R 5′-AGCTTATTCATCGCCACGTT -3′ | 33 |
| CTX-M Group-2 | F 5′-CGACGCTACCCCTGCTATT -3′  R 5′-CCAGCGTCAGATTTTTCAGG -3′ | 33 |
| CTX-M Group-8 | F 5′-TCGCGTTAAGCGGATGATGC -3′  R 5-AACCCACGATGTGGGTAGC -3′ | 33 |
| CTX-M Group-9 | F 5′-CAAAGAGAGTGCAACGGATG -3′  R 5′-ATTGGAAAGCGTTCATCACC -3′ | 33 |
| CTX-M Group 25/26 | F 5′-GCACGATGACATTCGGG -3′  R 5′-AACCCACGATGTGGGTAGC -3′ | 33 |

**Table S2: Distribution, prevalence of resistance genes and conjugation studies using *S. sciuri* isolated from poultry litter.**

| ***S. sciuri* isolate** | **Poultry farm** | **Antibiotic-resistant** | **Presence of gene** | **Presence of plasmid** | **Transconjugation** |
| --- | --- | --- | --- | --- | --- |
| ***S. sciuri* SCM1** | Farm A | Meropenem | - | - | NP |
| ***S. sciuri* SCM2** | Farm A | Meropenem | - | - | NP |
| ***S. sciuri* SCM3** | Farm A | Meropenem | *bla*_NDM-1_ | + | + |
| ***S. sciuri* SCM4** | Farm A | Meropenem | - | - | NP |
| ***S. sciuri* SCM5** | Farm A | Meropenem | *bla*_IMP_ | - | NP |
| ***S. sciuri* SCM6** | Farm A | Meropenem | - | - | NP |
| ***S. sciuri* SCM7** | Farm A | Meropenem | *bla*_NDM-1_ | - | NP |
| ***S. sciuri* SCM8** | Farm A | Meropenem | - | - | NP |
| ***S. sciuri* SCM9** | Farm A | Meropenem | - | - | NP |
| ***S. sciuri* SCM10** | Farm A | Meropenem | *bla*_IMP_ | - | NP |
| ***S. sciuri* SCM11** | Farm B | Meropenem | - | - | NP |
| ***S. sciuri* SCM12** | Farm B | Meropenem | - | - | NP |
| ***S. sciuri* SCM13** | Farm B | Meropenem | *bla*_IMP_ | - | NP |
| ***S. sciuri* SCM14** | Farm B | Meropenem | - | - | NP |
| ***S. sciuri* SCM15** | Farm B | Meropenem | *bla*_IMP_ | - | NP |
| ***S. sciuri* SCM16** | Farm B | Meropenem | - | - | NP |
| ***S. sciuri* SCM17** | Farm B | Meropenem | - | - | NP |
| ***S. sciuri* SCM18** | Farm B | Meropenem | *bla*_IMP_ | - | NP |
| ***S. sciuri* SCM19** | Farm C | Meropenem | *bla*_IMP_ | - | NP |
| ***S. sciuri* SCM20** | Farm C | Meropenem | - | - | NP |
| ***S. sciuri* SCM21** | Farm C | Meropenem | *bla*_IMP_ | - | NP |
| ***S. sciuri* SCM22** | Farm C | Meropenem | - | - | NP |
| ***S. sciuri* SCM23** | Farm C | Meropenem | - | - | NP |
| ***S. sciuri* SCM24** | Farm C | Meropenem | - | - | NP |
| ***S. sciuri* SCM25** | Farm C | Meropenem | *bla*_IMP_ | - | NP |
| ***S. sciuri* SCM26** | Farm C | Meropenem | - | - | NP |
| ***S. sciuri* SCC27** | Farm A | Cefotaxime | - | - | NP |
| ***S. sciuri* SCC28** | Farm A | Cefotaxime | - | - | NP |
| ***S. sciuri* SCC29** | Farm A | Cefotaxime | *bla*_CTX-M-9_ | - | NP |
| ***S. sciuri* SCC30** | Farm A | Cefotaxime | - | - | NP |
| ***S. sciuri* SCC31** | Farm A | Cefotaxime | *bla*_CTX-M-9_ | - | NP |
| ***S. sciuri* SCC32** | Farm A | Cefotaxime | - | - | NP |
| ***S. sciuri* SCC33** | Farm A | Cefotaxime | - | - | NP |
| ***S. sciuri* SCC34** | Farm A | Cefotaxime | - | - | NP |
| ***S. sciuri* SCC35** | Farm A | Cefotaxime | *bla*_CTX-M-9_ | - | NP |
| ***S. sciuri* SCC36** | Farm B | Cefotaxime | - | - | NP |
| ***S. sciuri* SCC37** | Farm B | Cefotaxime | *bla*_CTX-M-2_ | - | NP |
| ***S. sciuri* SCC38** | Farm B | Cefotaxime | - | - | NP |
| ***S. sciuri* SCC39** | Farm B | Cefotaxime | *bla*_CTX-M-9_ | - | NP |
| ***S. sciuri* SCC40** | Farm B | Cefotaxime | - | - | NP |
| ***S. sciuri* SCC41** | Farm B | Cefotaxime | - | - | NP |
| ***S. sciuri* SCC42** | Farm B | Cefotaxime | - | - | NP |
| ***S. sciuri* SCC43** | Farm C | Cefotaxime | - | - | NP |
| ***S. sciuri* SCC44** | Farm C | Cefotaxime | *bla*_CTX-M-9_ | - | NP |
| ***S. sciuri* SCC45** | Farm C | Cefotaxime | - | - | NP |
| ***S. sciuri* SCC46** | Farm C | Cefotaxime | - | - | NP |
| ***S. sciuri* SCC47** | Farm C | Cefotaxime | - | - | NP |
| ***S. sciuri* SCC48** | Farm C | Cefotaxime | - | - | NP |
| ***S. sciuri* SCC49** | Farm C | Cefotaxime | - | - | NP |
| ***S. sciuri* SCC50** | Farm C | Cefotaxime | - | - | NP |

‘+’ – positive isolate; ‘-’ – negative isolate; NP – not performed
